# Metagenomic and transcriptomic signatures of periodontitis in companion dogs

**DOI:** 10.64898/2026.03.31.715430

**Authors:** Alex Grier, Jennifer K. Grenier, Michael J. Byron, Nadine Fiani, Nicole Traver, Alex M. Valm, Santiago Peralta

## Abstract

**Background:** Periodontitis (Perio) is a progressive oral disease characterized by inflammation and degradation of the periodontal apparatus and is associated with local and systemic morbidity including loss of teeth, cardiovascular disease, and diabetes mellitus, among others. Perio is highly prevalent in domestic canines and exhibits certain parallels in pathogenesis and pathophysiology to Perio in humans, although standard treatments are less effective. In both species, a complex interplay between oral microbiota and host immune response is implicated in the etiology of Perio but is not fully understood.

**Results:** Using shotgun metagenomics and RNA-seq on oral samples from companion dogs, we identify features of the oral microbiome and host transcriptional profile that are associated with Perio and its progression. We observe differences in microbiota composition between Perio and non-Perio animals that are largely consistent with what has been described in humans but also identify several species that are distinctly associated with canine Perio. We observe an abrupt shift in host gene expression related to immune response and tissue structure that is associated with disease severity, specifically the progression from mild periodontal disease (PD) to more severe Perio and the initiation of clinical attachment loss. The gingival plaque microbiota exhibits a parallel dynamic, with distinct compositional profiles in mild, moderate, and severe PD. We then examine several of the known mechanistic components of the keystone pathogen hypothesis of PD, identifying specific commonalities between canine and human pathologies, including the involvement of *Porphyromonas* species and related virulence factors. Additionally, we show infiltration of gingival tissue by *Porphyromonas* and *Tannerella* spp. via fluorescence microscopy. Finally, we assess correlations between host gene expression and microbial metabolic pathways which suggest additional potential virulence factors.

**Conclusions:** This work elucidates the metagenomic and transcriptomic signatures of Perio in companion dogs with the goals of informing veterinary medicine, evaluating the potential of canines as a model organism for the study of Perio, and clarifying the relationship between Perio development and progression, the oral microbiota, and the localized host response. Our findings provide insight into the etiopathogenesis of canine Perio and its relationship to human Perio and suggest novel targets of potential translational interest.

## Background

Periodontal disease (PD) is clinically characterized by gingival inflammation (i.e., gingivitis), that is often accompanied by irreversible clinical attachment loss due to progressive periodontal tissue destruction (i.e., periodontitis) (Martínez-García, 2021). As in humans and other mammalian species, periodontal disease is highly prevalent in companion dogs (Harvey, 1998). Untreated or unresponsive PD is associated with local and systemic morbidity including loss of teeth, pathological jaw fracture, oronasal and oroantral fistula formation, cardiovascular disease, diabetes mellitus, among others (Hajishengallis G., 2022). For decades, standard-of-care preventative, therapeutic and maintenance strategies have revolved around mechanical removal of subgingival plaque and calculus deposits (Matthews, 2014). However, response rates are variable, and susceptible individuals are frequently refractory to therapy, underscoring the need to develop novel interventions (Kinane, 2017). Understanding the etiopathogenetic mechanisms of disease is essential to formulate predictable, rational, and more effective clinical solutions.

Historical etiopathogenetic models of PD include the specific plaque hypothesis (Loesche, 1976), which attributes a pathogenic role to certain bacterial species; the non-specific plaque hypothesis, which proposes that undisturbed subgingival plaque accumulation regardless of its microbial composition is sufficient to induce disease (Theilade, 1986); and the ecological plaque hypothesis, whereby disease is determined by compositional shifts in the subgingival microbial communities due to changing microenvironmental conditions (Marsh, 1994). Of note, these models were developed using culture-based methodologies or targeted molecular techniques, and don’t necessarily align with the natural course and clinical spectrum of disease (Belibasakis, 2023). More recently, unbiased exploration of the subgingival microbial communities and mechanisms of local host response using next-generation sequencing and other high-throughput approaches, have uncovered additional mechanistic complexity leading to new proposed models including the ‘keystone pathogen’ hypothesis (Hajishengallis G. D., 2012). The latter assigns e.g., *Porphyromonas gingivalis*, even when present in relatively low numbers, the ability to induce a local inflammatory host response that favors the transition of the subgingival microbiota from a symbiotic to a dysbiotic state that further promotes inflammation, thus representing a vicious pathological cycle (Olsen, 2017).

Much of what is currently known about the mechanisms of PD has been established using laboratory animals including non-human primates, rats, mice, and dogs (Oz, 2011). Interestingly, although animal models are useful for testing biologically relevant hypotheses, they are not ideal for pre-clinical applications (Frangogiannis, 2022).

Conversely, given the clinical, immunological, pharmacological, genetic, and environmental parallels that exist between companion animals and people with spontaneous disease, they are generally considered a more robust model system for translational applications (Kol, 2015). In this context, better understanding the etiopathogenetic mechanisms of PD in companion dogs may help to leverage and validate them as a potentially useful pre-clinical model of corresponding disease in people while simultaneously informing candidate approaches of veterinary value (Albuquerque, 2012). Therefore, the aims of this study were to characterize the subgingival microbiota and host transcriptional profile of naturally occurring PD in companion dogs using a shotgun metagenomics and bulk RNA-seq approach to gain mechanistic insights of potential comparative, translational and veterinary relevance.

## Materials and Methods

### Clinical Samples

Subgingival plaque samples were collected from adult and otherwise systemically healthy client-owned dogs of varied age, sex, and breed presented to the Dentistry and Oral Surgery Service at the Cornell University Hospital for Animals, with no recent history (i.e., 4 weeks or less) of topical or systemic antibiotic or immunosuppressive drug administration. Signed dog owner written consent was obtained prior to sample collection. Samples were collected sterilely using an endodontic absorbent paper point, as previously described (Rodrigues M. X., 2019) (Rodrigues M. X., 2021), while dogs were under general anesthesia undergoing standard-of-care procedures. Sampling occurred after full-mouth radiographs had been obtained and prior to any chemical or mechanical, topical, or systemic interventions that could alter the subgingival microenvironment. From each dog, the most severely periodontally diseased sites were prioritized; preference was given to canine and carnassial teeth when applicable. Each sample was assigned a PD severity score based on radiographic (i.e., percentage of alveolar bone loss) and clinical findings (i.e., gingival index, and clinical attachment loss based on probing depth and gingival recession determined after sampling). Based on severity score, the three PD categories were defined: mild, defined as teeth with some degree of gingivitis but no clinical attachment loss; moderate, defined as teeth with clinical attachment loss of less than 50%; and severe, defined as teeth with more that 50% of clinical attachment loss. Samples with mild PD were designated as non-periodontitis and moderate and severe samples were designated as periodontitis (Perio). Samples were immediately placed in a sterile Eppendorf tube and stored in -80 C until analyzed. Matching gingival margin tissue samples measuring approximately 2 mm^3^ were collected from a subset of dogs during standard-of-care surgical dental extraction. Tissue samples were immediately flash-frozen and stored in liquid nitrogen until analyzed. From some dogs, immediately adjacent tissue samples were preserved in formalin and embedded in paraffin for sectioning. Pertinent metadata was tabulated from each dog including breed, body weight, sex, and age. All sampling procedures, group assignment and clinical data collection were performed or directly supervised by a board-certified veterinary dentist (SP) and complying with protocols that had been previously reviewed and approved by Cornell University’s Institutional Animal Care and Use Committee (protocols #2015-0117 and #2015-0219).

### DNA Extraction and Sequencing

Endodontic absorbent paper point samples were taken from sterile Eppendorf tubes and transferred into a Qiagen PowerBead glass 0.1 mm tube (13118-50). Using a Promega Maxwell RSC PureFood GMO and Authentication Kit (AS1600), 1mL of CTAB buffer & 20μl of RNAse A Solution was added to the PowerBead tube containing the sample.

The sample/buffer was mixed for 10 seconds on a Vortex Genie2 and then incubated at 95°C for 5 minutes on an Eppendorf ThermoMixer F2.0, shaking at 1500 rpm. The tube was removed and clipped to a horizontal microtube attachment on a Vortex Genie2 (SI-H524) and vortexed at high-speed for 20 minutes. The sample was removed from the Vortex and centrifuged on an Eppendorf Centrifuge 5430R at 40°C, 12700 rpm for 10 minutes. Upon completion, the sample was centrifuged again for an additional 10 minutes to eliminate foam. Any remaining foam and particulates were carefully removed using P1000 pipette. DNA extraction was then processed via the Promega MaxPrep Liquid Handler and Promega Maxwell RSC 48 instruments. DNA was quantified using Quant-iT dsDNA High Sensitivity Assay Kit using Promega GloMax plate reader on a microplate (655087). Library generation followed the Illumina Nextera XT DNA Library Prep Kit Reference Guide: https://support.illumina.com/content/dam/illumina-support/documents/documentation/chemistry_documentation/samplepreps_nextera/nextera-xt/nextera-xt-library-prep-reference-guide-15031942-05.pdf

DNA sequencing libraries were washed using Beckman Coulter AMPure XP magnetic beads. Library quality & size verification was performed using PerkinElmer LabChip GXII instrument with DNA 1K Reagent Kit (CLS760673). Library concentrations were quantified using Quant-iT dsDNA High Sensitivity Assay Kit using Promega GloMax plate reader on a microplate (655087). Library molarity was calculated based on library peak size & concentration. Libraries were normalized to 2nM using the PerkinElmer Zephyr G3 NGS Workstation (133750) and pooled together using the same volume across all normalized libraries into a 1.5ml Eppendorf DNA tube (022431021). Pooled libraries were sequenced on the Illumina HiSeqX instrument at loading concentration of 10pM, paired-end 150bp.

### RNA Extraction and Sequencing

For RNA isolation, frozen host tissues (∼1 gm) were homogenized in 1 mL of Trizol (Thermo Fisher) using 2.8 mm ceramic beads (Hard Tissue Homogenizing Mix, VWR). RNA extraction was performed with a modified Trizol method as follows: after the addition of chloroform and phase separation of the Trizol lysate, the aqueous phase was combined with an equal volume of 100% ethanol and loaded onto a Zymo-Spin column and purified using the Quick-RNA Prep Kit (Zymo Research). For all samples, RNA concentration was measured with a Nanodrop (Thermo Fisher), and integrity was determined with a Fragment Analyzer (Agilent). Ribosomal RNA was depleted with the NEBNext rRNA Depletion Kit v2 (Human/Mouse/Rat; New England Biolabs) using 500 ng input total RNA. All RNA-seq libraries were generated with the NEBNext Ultra II Directional library prep kit (New England Biolabs) and at least 40M 2x150bp paired-end reads were generated for each sample on a NovaSeqX instrument (Illumina).

### Data analysis

For shotgun metagenomics, quality-based filtering and removal of host sequences and adapter and quality trimming was performed using KneadData (https://huttenhower.sph.harvard.edu/kneaddata/) version 0.10.0, using the GCF_000002285.5 canine reference genome. Taxonomic composition was calculated using MetaPhlAn4 (Blanco-Míguez, 2023) with the mpa_v30_CHOCOPhlAn_201901 reference database. Relative abundances of UniRef90 gene families and metabolic pathways were quantified using HUMAnN3 v3.6 (Beghini, 2021). Gene families and pathways were renormalized to relative abundances after removing unmapped/ungrouped counts.

Relative abundances of taxa, pathways, and gene families were converted to integers by multiplying by the number of post-filtering reads in each sample and rounding. Using these integer counts, alpha and beta diversity were calculated using QIIME2 (Bolyen, 2019). Group differences in alpha diversity were calculated using the Wilcoxon rank-sum test. Group differences in beta diversity were calculated using PERMANOVA and visualized using Principal Coordinate Analysis values generated by QIIME2. Differential abundance analysis of taxa and pathways were performed using Maaslin3 (Nickols, 2026) with TSS normalization, LOG transformation, augment and standardize set to true, max_significance set to 0.1, median_comparison_abundance set to true, median_comparison_prevalence set to false, min_prevalence set to 0.1, min_abundance set to 0.0, and otherwise default parameters. Unbiased clustering of metagenomic samples based on taxonomic composition was performed using the DirichletMultinomial R Package version 1.50.0 (Holmes, 2012), based on species present in at least ten samples. Correlation analysis between microbial pathways and gene expression was performed using SECOM version 2.10.1 (Lin, 2022) including only genes that were differentially expressed between clades (see Results) or disease groups and pathways that were differentially expressed between disease groups using the following parameters: pseudo = 0, prv_cut = 0.25, lib_cut = 1000, corr_cut = 0.5, wins_quant = c(0.05, 0.95), R = 1000, thresh_hard = 0.3, max_p = 0.005, all other default. RNA-Seq differential expression was performed using DESeq2 (Love, 2014).

Over-representation analysis of differentially expressed genes labeled as their human homologs was performed using the PANTHER version 19.0 (Mi, 2019) web service with the PANTHER GO-Slim database or the Reactome database (Milacic, 2024), using the human genome as the background reference.

For RNA-seq data, raw reads were trimmed for low-quality and adaptor sequences and filtered for minimum length with TrimGalore v0.6.6 (http://www.bioinformatics.babraham.ac.uk/projects/trim_galore/), a wrapper for cutadapt (Martin, 2011) and fastQC (http://www.bioinformatics.babraham.ac.uk/projects/fastqc/) using parameters ‘--nextseq-trim=20 -O 1 -a AGATCGGAAGAGC --length 50 --fastqc’. Trimmed reads were mapped to the reference genome/transcriptome (Ensembl CanFam3.1) with STAR v2.7 (Dobin, 2013) using these parameters: ‘--outSAMstrandField intronMotif, --outFilterIntronMotifs RemoveNoncanonical, --outSAMtype BAM SortedByCoordinate, --outReadsUnmapped Fastx and --quantMode GeneCounts’, which also generated raw count outputs per annotated gene.

Sample clustering and differential gene expression were analyzed with SARTools^51^ and DESeq2 using these parameters: ‘fitType parametric, cooksCutoff TRUE, independentFiltering TRUE, alpha 0.05, pAdjustMethod BH, typeTrans VST, and locfunc median’. Canine gene symbols were converted to human gene symbols using Biomart (Ensembl) one-to-one orthology assignments to enable analysis with gene sets in MSigDB (Liberzon, 2015). The human ortholog gene symbols and log2-fold-change values for expressed genes (at least one group with average normalized counts > 50) were used for GSEA (Subramanian, 2005) ‘Preranked’ analysis.

### Fluorescent In Situ Hybridization

Genus-specific encoding probes were designed using the probe design software tool Decipher with the Canine Oral Microbiome Database (Wright, 2014) (Dewhirst, 2012). Three probes were designed for *Porphyromonas* to provide better species coverage, and one probe for *Tannerella*. The *Porphyromonas* probes: POR1246, POR285, and POR595, have the following sequences, respectively: CCTGTCGCCAGGTAGCTGC, TCTCAGTTCCCCTACCCA, and CTGCAGCTTTTCACCACTGACTT. The *Tannerella* probe (TAN167) sequence is: CGGGACCCCTGTTTTATG. The previously described EUB338 probe: GCTGCCTCCCGTAGGAGT was used to label all bacteria (Amann, 1990). Readout probes were conjugated to Alexa Fluor-514 for *Porphyromonas*, Alexa Fluor-555 for *Tannerella* and Alexa Fluor-594 for EUB (ThermoFisher). FISH labeling was carried out as previously described (Valm, 2011). After FISH labeling, tissues were stained with DAPI (0.55 µM) for 10 min. Samples were dried in an ethanol series then mounted in ProLong Gold antifade reagent (ThermoFisher).

### Image Acquisition and Processing

Spectral images were acquired using a Zeiss LSM980 laser scanning confocal microscope with the 405 nm, 488 nm, 561 nm, and 639 nm laser lines. Images were obtained as z-stacks between 8-30 slices using a 20x/0.8 NA objective (low magnification, large field of view) or 63x/1.4 NA objective (high magnification, small field of view). Large field of view images were acquired as tile scans. Reference spectra for linear unmixing were obtained from images of *E. coli* (ATCC 10798) that had been labelled with the same fluorophores used for taxon-specific probes and acquired with the same settings as the tissue samples. Linear unmixing and image stitching were performed in Zeiss Zen Blue Software. After unmixing, the images were processed in Fiji (Schindelin, 2012). Brightness and contrast were adjusted uniformly across the entire images and z stacks were transformed into maximum intensity projections.

## Results

A total of 101 client-owned adult dogs participated in this study. The average age of dogs was 7.9±3.6 years, the average body weight was 14.2±12.1 kg, and the sex distribution was 53.5% males (52 castrated and 2 intact) and 46.5% females (41 spayed and 1 intact), representing a total of 39 pure or mixed breeds. A single subgingival plaque sample from each subject underwent shotgun metagenomic sequencing and a subset of 22 subjects were selected for RNA-seq and FISH experiments using gingival tissue samples. From each dog, the most severely periodontally diseased sites were prioritized; preference was given to canine and carnassial teeth when applicable. Each sample was assigned a disease severity score from 0 to 4 based on radiographic and clinical findings. The median and mode of severity scores was 3.

Metagenomic sequencing yielded a total of 981M read pairs. The mean percentage of non-canine sequences in the metagenomic samples was 15.8% (+/-14.4%), corresponding to a mean read count of 2.6M (+/- 2.5M) after QC filtering and canine sequence removal. Eight low read-count (post-filtering) samples were excluded from metagenomic analysis due to having no identifiable taxa or metabolic pathways. In the remaining samples, a total of 124 unique species were identified. For the purposes of preliminary analysis, disease was partitioned into two groups based on severity: scores of 0-1 corresponded to teeth with some degree of gingivitis but no clinical attachment loss and were labeled “non-periodontitis” (non-Perio; n = 29) while scores >= 2 were labeled “periodontitis” (Perio; n = 64). We began with an unbiased characterization of the differences between groups. Alpha diversity, as measured by both richness and the Shannon index, was elevated in the Perio group (Figure 1A; p = 0.0005 and p = 0.0083, respectively). Beta diversity, as measured by Bray-Curtis dissimilarity, was significantly different between non-Perio and Perio groups (Figure 1B; PERMANOVA p = 0.003). We observed 18 species that differed in either prevalence or abundance between Perio and non-Perio using a significance threshold of FDR <= 0.1 (Figure 1C). *Parvimonas micra*, *Bergeyella zoohelcum*, *Capnocytophaga canis*, and *Capnocytophaga canimorsus* were elevated in the non-Perio group. *Bacteroides pyogenes*, *Campylobacter rectus*, *Treponema denticola*, *Bacteroides heparinolyticus*, *Bulleidia extructa*, *Porphyromonas crevioricanis*, *Campylobacter showae*, *Tannerella forsythia*, *Filifactor alocis*, *Fretibacterium fastidiosum*, *Peptostreptococcaceae bacterium oral taxon 113*, *Methanobrevibacter oralis*, *Desulfomicrobium orale*, and *Treponema socranskii* were elevated in the Perio group. Using the same significance threshold, 101 microbial metabolic pathways were identified as being differentially abundant between groups, the top 25 most significant of which are shown (Figure 1E; Supplementary Table 1).

**Fig. 1.**
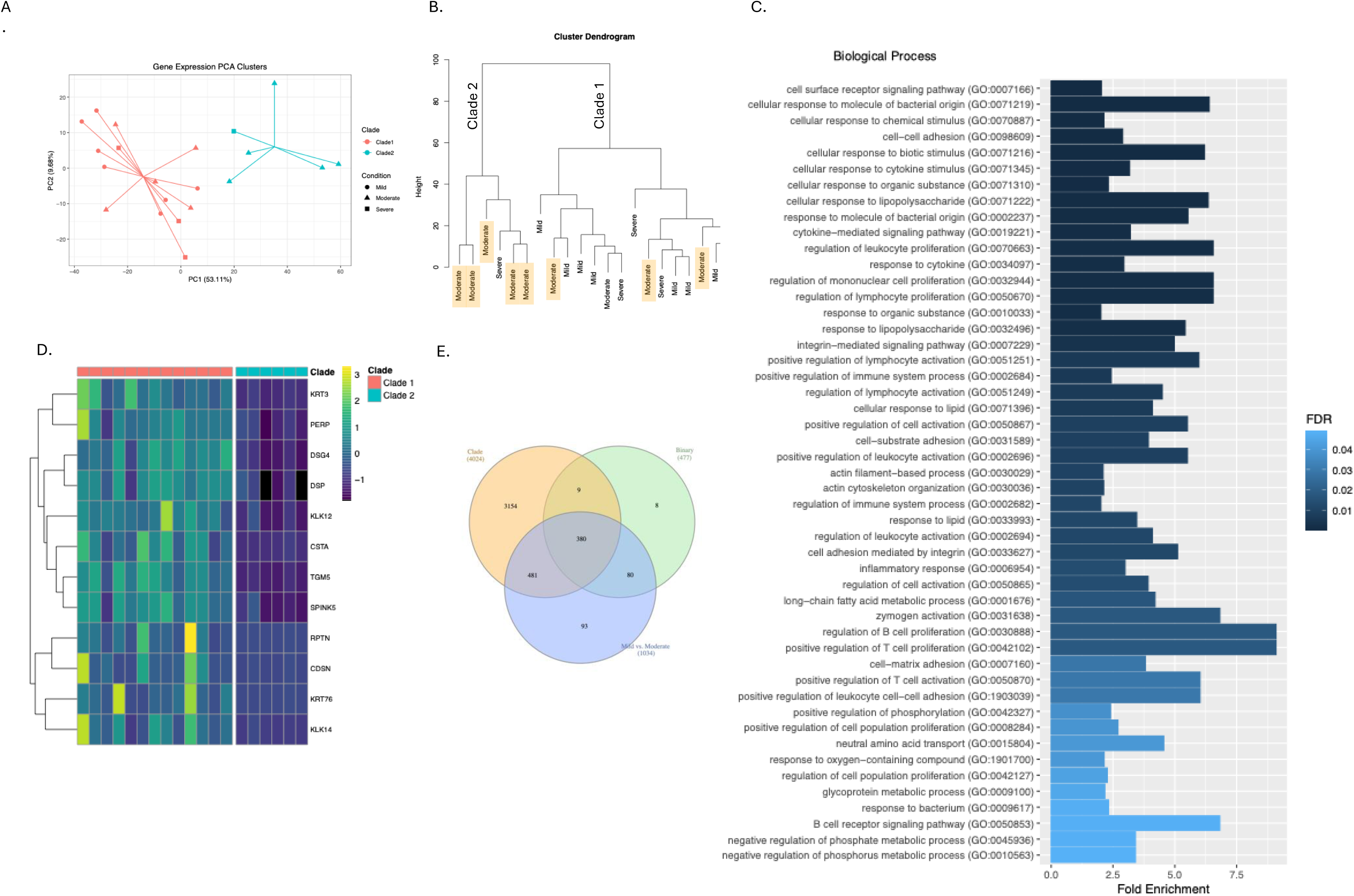
Characterization of periodontitis vs. non-periodiontitis. A) Alpha diversity differences between groups. B) Beta diversity differences between groups. C) Heatmap of the scaled relative abundances of differentially abundant species (FDR <= 0.1) between groups. D) Heatmap of the scaled expression of genes from the Davanian periodontitis gene set that were differentially abundant between groups in this data set. E) The top 25 most significantly differentially abundant microbial metabolic pathways between groups, colored by FDR with the coefficient plotted on the x-axis. F) Over-represented biological processes based on the set of differentially abundant genes between groups.

RNA-seq yielded mapping rates exceeding 80% of reads and ∼15M reads mapping to annotated genes for each sample. One sample was discarded due to being an extreme outlier by PCA analysis. Of the remaining 21 samples, 7 were from the non-Perio group and 14 were from the Perio group. Reads were initially mapped to the canine reference genome and genes were subsequently relabeled with their human homologs when possible. All subsequent analysis refers to the human homolog labels and excludes genes without a 1-to-1 human ortholog. Differential expression analysis between the non-Perio/Perio groups identified 477 differentially expressed genes using a significance threshold of FDR <= 0.1. To assess potential parallels between canine and human periodontitis, we performed Gene Set Enrichment Analysis against the set of human periodontitis-associated genes identified in (Davanian, 2012), revealing highly significant enrichment (p ≈ 0.0, normalized enrichment score = 8.84, Supplementary Figure 1). To guide preliminary interpretation, we visualized the expression of the genes from the Davanian set that were significantly differentially abundant in our dataset (Figure 1D). To characterize high-level functional differences in gene expression between groups, we used our entire list of differentially expressed genes with the human genome as background reference to perform overrepresentation analysis against the PANTHER GO-Slim database of biological processes, filtering for processes significantly enriched with an FDR <= 0.05 and an enrichment of >= 2.0 fold (Figure 1F). Processes related to cellular motility (12/24) and immune response (4/24) comprised the majority of these results.

We next investigated a more fine-grained resolution of disease severity, binning subjects into one of three categories based on PD severity score: mild (score 0-1), moderate (score 2-3), and severe (score 4). In the course of exploratory analyses, we observed that in PCA plots of the RNA-seq data, two distinct clades emerged (Figure 2A & 2B). Examination of sample severity group assignments revealed that one clade (labeled as Clade 2) was comprised almost entirely of moderate severity subjects (Figure 2B); five of the six samples in Clade 2 came from moderate severity subjects and one from a subject with severe disease. To further interrogate the differences between clades, we performed differential expression analysis and identified 4024 genes with FDR <=0.1. For preliminary functional characterization of the differences between clades, we used the subset of 986 highly significant (FDR <= 0.001) genes for overrepresentation analysis, identifying 49 overrepresented biological processes with FDR <=0.05 and >=2.0-fold enrichment (Figure 2C). These processes were primarily related to a shift in immune response (19/49 significant processes), proliferation and adhesion (14/49), and cell signaling (9/49), consistent with increased inflammation and degradation and disruption of tissue structure/detachment. In order to try to put a finer point on the functional differences between the clades, we then performed overrepresentation analysis using a smaller subset of 258 extremely significant genes (FDR <= 0.00001) against the Reactome pathway database. This yielded a single significant pathway, Keratinization, which was overrepresented with 4.31-fold enrichment and an FDR of 0.0331. Expression of the extremely significantly differentially expressed genes represented in the Reactome Keratinization pathway was consistently down-regulated (Figure 2D). To contextualize the scope of the differences between clades relative to differences between disease severity groups, we constructed a Venn diagram of the differentially expressed genes resulting from clade-clade comparison, non-Perio vs. Perio comparison (Binary), and the mild vs. moderate severity group comparison (Figure 2E). This revealed that far more genes were differentially expressed between the clades than between the disease/severity groups. Interestingly, there were no significantly differentially expressed genes between the severe and mild disease groups using an FDR threshold of <= 0.1. Similarly, there were no significantly differentially expressed genes between the severe and moderate groups.

**Fig 2.**
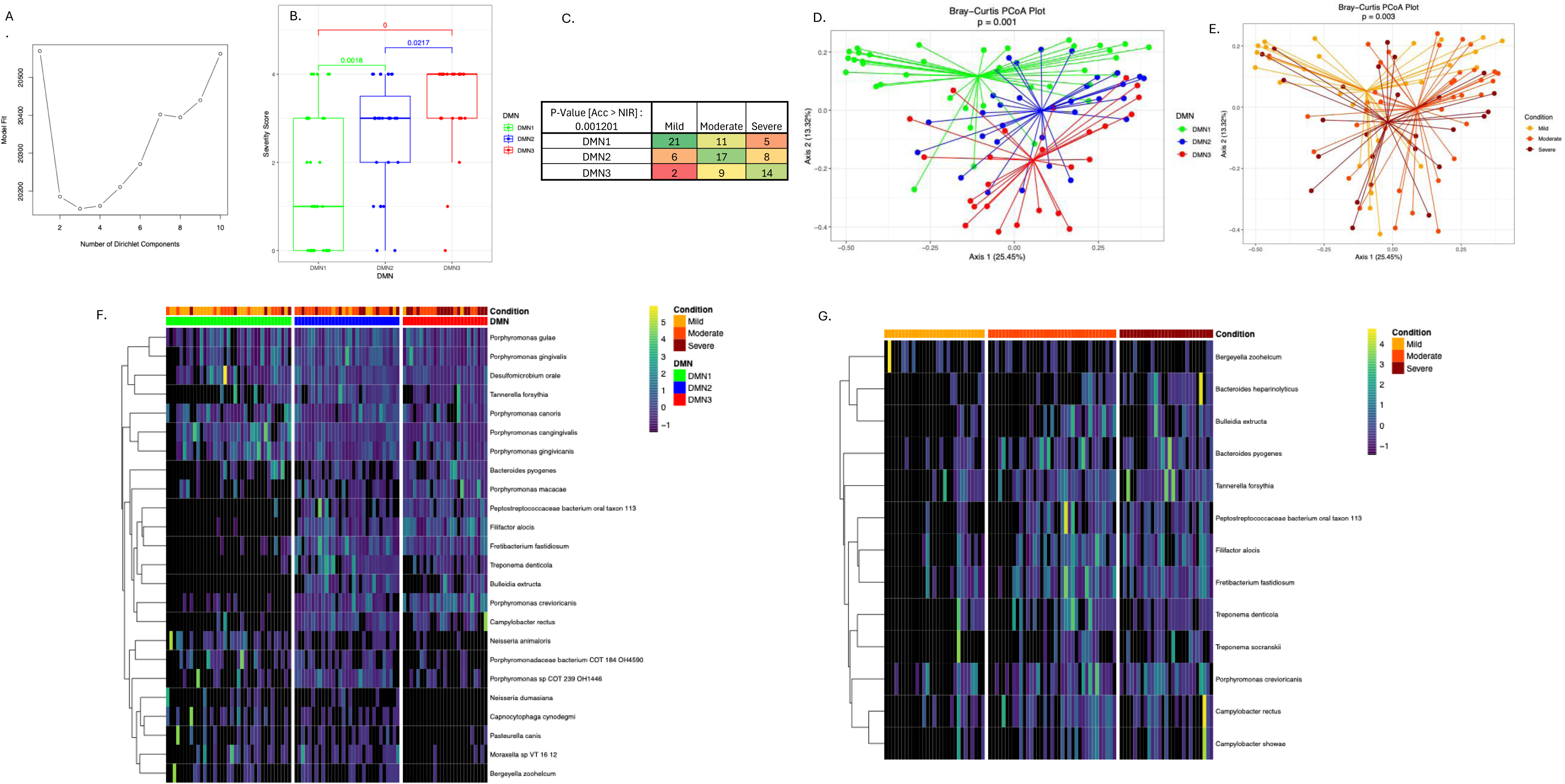
Clades based on host gene expression profiles. A) PCA of RNA-Seq samples, colored by clade. B) Dendrogram of samples labeled by severity group, using PC1-PC3 distances. C) Over-represented processes based on highly significantly (FDR <= 0.001) differentially abundant genes between clades. D) Heatmap of the scaled expression of differentially abundant genes between clades within the Keratinization pathway. E) A Venn diagram of the differentially expressed genes resulting from clade-clade comparison, non-Pr vs. Pr comparison (Binary), and the mild vs. moderate severity group comparison.

We then sought to assess whether the overall composition of microbiota corresponded to disease severity. We first performed unsupervised clustering of the metagenomic samples based on species-level taxonomic composition using a Dirichlet Multinomial Model. We repeatedly fit the model with the number of clusters set to one through ten and assessed goodness-of-fit using the Laplace score, which identified the three-cluster model as the best fit to the data (Figure 3A). These unsupervised clusters correspond well to disease severity score and severity groups (Figure 3B & 3C, p = 0.001). Visualizing these clusters in PCoA space based on Bray-Curtis dissimilarity revealed a more significant difference than that observed between severity groups (Figure 3D & 3E). Using the posterior mean difference between our fitted three-cluster model and a single-component null model, we identified the species primarily driving the clustering (Figure 3F shows the species accounting for 90% of the difference compared to the null/single-component model). Interestingly, just seven species accounted for over 50% of the difference (in descending order): *Porphyromonas gulae*, *Porphyromonas crevioricanis*, *Porphyromonas cangingivalis*, *Porphyromonas macacae*, *Porphyromonas gingivicanis*, *Filifactor alocis*, and *Porphyromonas gingivalis*.

**Fig. 3.**
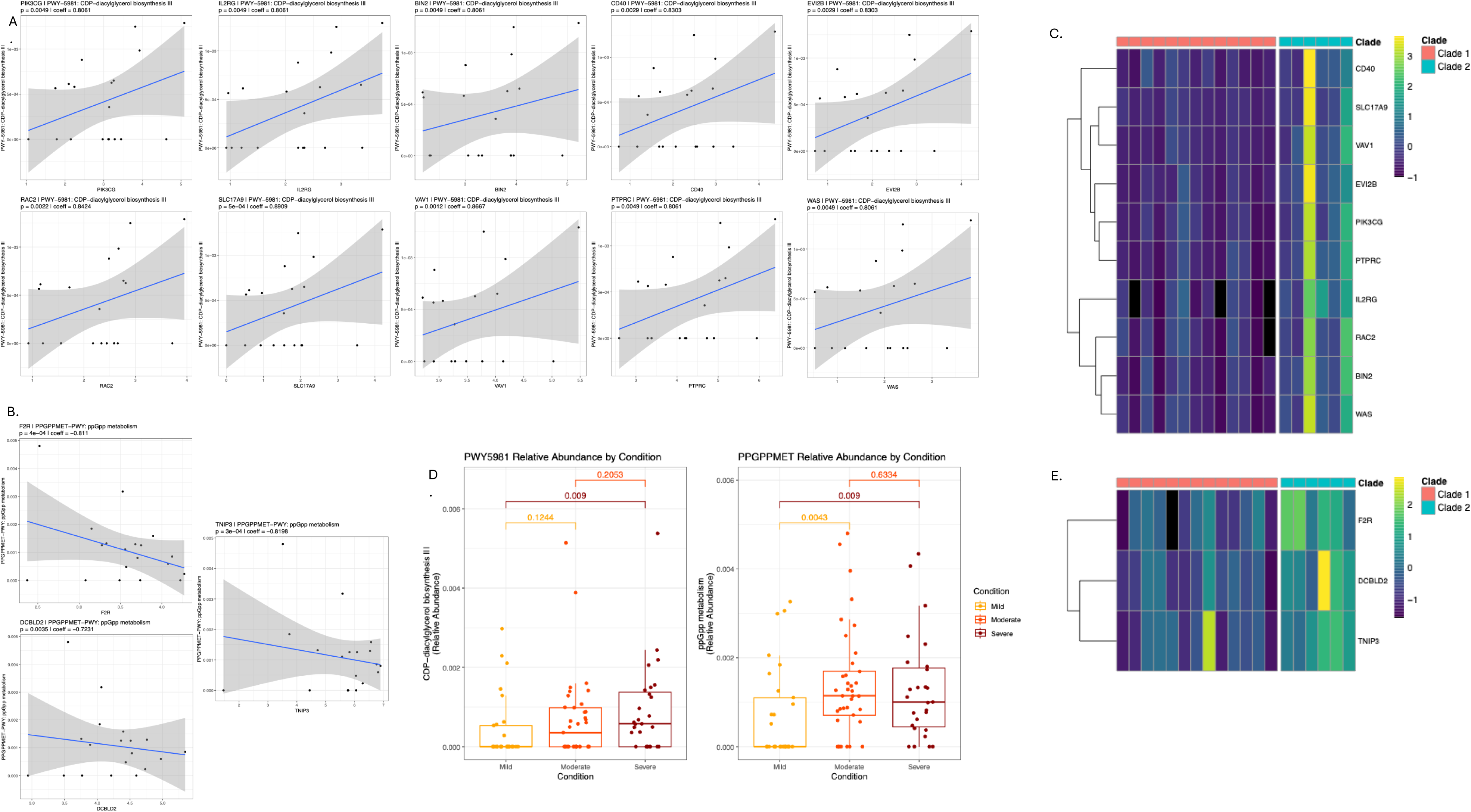
Unbiased clustering of microbiota samples and differences based on disease severity group. A) Laplace scores of model fit for models with 1 through 10 clusters. B). Distribution of disease severity scores with DMN clusters. C) Distribution of severity groups across DMN clusters and significance test of accuracy (Acc) compared to No Information Rate (NIR). D) Bray-Curtis PCoA and PERMANOVA based on the DMN clusters. E) Bray-Curtis PCoA and PERMANOVA based on the severity groups. F) Heatmap of the scaled relative abundance of the species accounting for 90% of the difference compared to the null/single-component model. G) Heatmap of the scaled relative abundance of the differentially abundant species between severity groups (FDR <= 0.1).

Additionally, fluorescence microscopy of gingival tissue samples showed infiltration by *Porphyromonas spp.* and *Tannerella spp*. (recall that *Tannerella forsythia* was significantly elevated in Perio vs. non-Perio samples). To assess the overlap between species driving the unsupervised clustering and species distinguishing the severity groups, we performed differential abundance and differential prevalence analysis on the severity groups, identifying 13 species that differed significantly with an FDR <= 0.1 (Figure 3G). Of these, 10 were among the top drivers of the unsupervised clusters, accounting for 34% of the difference from the null model.

Having characterized the microbiota and gene expression of Perio in an unbiased manner, we then examined biomarkers related to known mechanisms of disease progression in humans, namely complement-dependent dysbiotic inflammation as described by the keystone pathogen hypothesis. Based on our unbiased observation of two clades of samples with very different host gene expression profiles, and given that one of the two clades was comprised almost exclusively of moderate disease, we sought to investigate whether that group of samples exhibited a distinct signature of disease onset or stepwise progression. We therefore began by looking at C5A complement receptor expression in these two sets of samples (Figure 4A). Not only did we find that *C5AR1* expression was dramatically increased in Clade 2/Moderate Clade (p = 0.0003), but we observed that the only Clade 2 sample which was *not* in a moderate disease state (it was instead in a severe disease state) had notably lower *C5AR1* expression than the other samples in the clade (Figure 4A, circled in red). That one sample had *C5AR1* expression consistent with the 3^rd^ quartile of Clade 1 samples, while all other Clade 2 samples (all moderate disease) had higher *C5AR1* expression than *any* of the Clade 1 samples. Given that complement-dependent dysbiotic inflammation in human PD is attributed to the effects of bacterially-produced gingipains, we then searched for gingipain homolog sequences in the metagenomic samples and identified five such genes attributed to four *Porphyromonas* species (UniRef90_P95493 and UniRef90_A0A1L5IZD9 to *P. gingivalis* and *P. gulae*; UniRef90_A0A0A2EZF3 to *P. gulae* only; UniRef90_S4PFY5 to *P. crevioricanis*, and UniRef90_A0A0A2G4V4 to *P. gingivicanis*) which were significantly increased in prevalence in moderate or severe disease with FDR <= 0.1 (Figure 4B). While there is definitive evidence of enzymatically active gingipains in *P. gingivalis* and *P. gulae*, the status of these genes in other *Porphyromonas* species is currently inconclusive (Lenzo, 2016). With the data available, we are only able to show that sequences homologous to canonical gingipains align to the reference genomes of other *Porphyromonas* species, and that the abundance of those sequences are highly correlated with the abundance of these species (Figure 4C) – our data is insufficient to show that other *Porphyromonas* species produce functional gingipains. Because we suspected that the onset of gingipains-induced complement-dependent dysbiotic inflammation was the primary feature of the Clade 2 samples that distinguished their host gene expression profiles from other samples, we looked at the expression of other known gingipains-responsive genes and other components of the complement system. We found a number of these genes to be significantly upregulated (FDR <= 0.1) in Clade 2 including Toll-like receptors 2 (*TLR2*) and 6 (*TLR6*), Factor 2 receptor (*F2R*), C3A receptor (*C3AR1*), transferrin (*TF*), Adhesion Molecule 1 (*ICAM1*), *AKT3*, and Insulin Receptor protein (*INSR*) (Figure 4D). Given that *Porphyromonas* species were the primary drivers of our unsupervised clusters of microbiota samples, we examined the distribution of sequence homologs of the canonical P*. gingivalis gingipains* rgpB (UniRef90_P95493) across clusters and severity groups, revealing that they were significantly elevated in the moderate disease samples (p = 0.007) and in the cluster corresponding to moderate severity relative to the other two clusters (Figure 4E). Finally, we examined the abundance of sequence homologs of the key *P. gingivalis* virulence factor fimA across unsupervised clusters and severity groups and found it to be significantly elevated the cluster corresponding to severe disease (p = 0.03) and in moderate and severe groups (Figure 4F).

**Fig 4.**
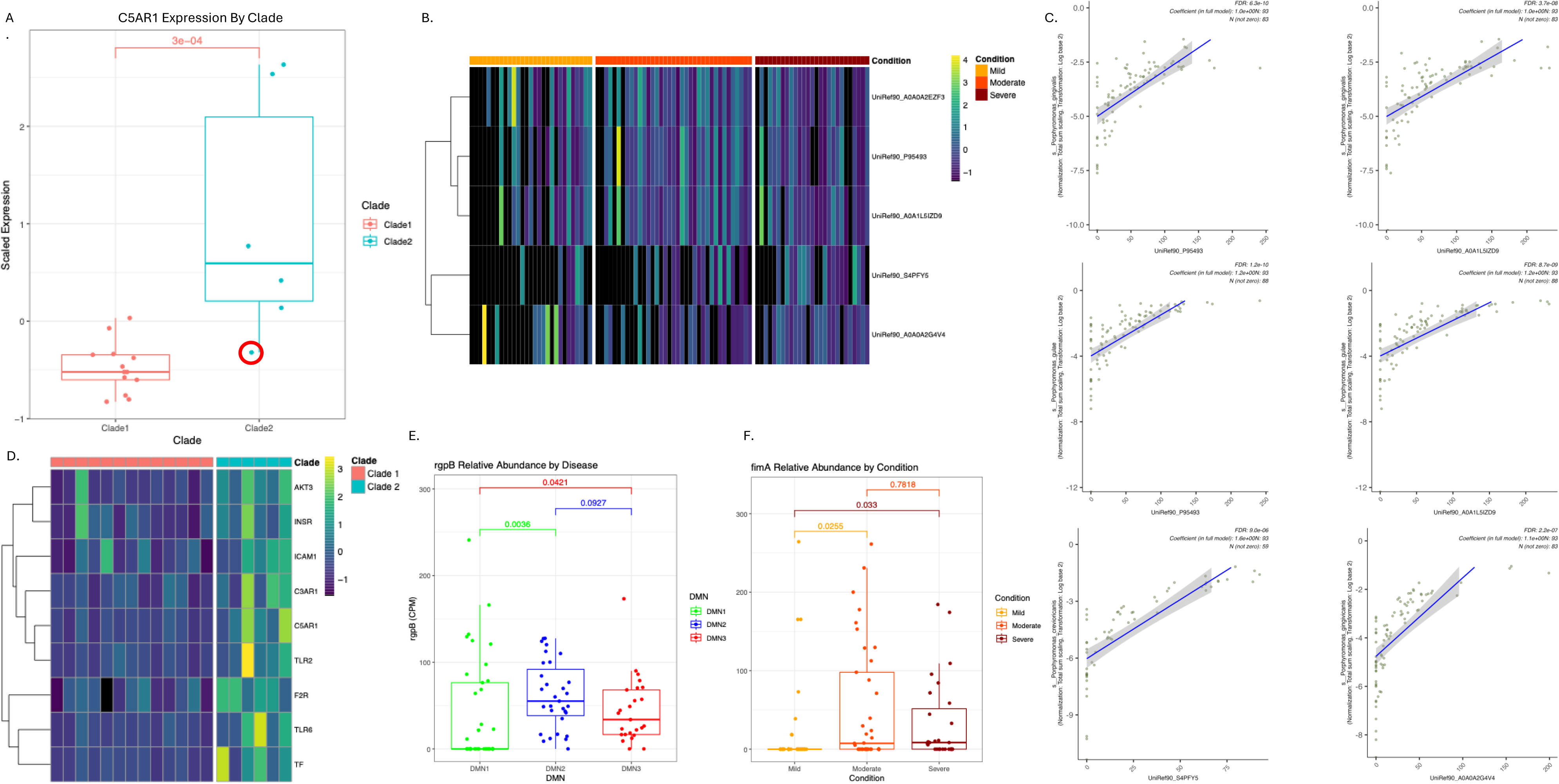
Gingipains, their targets, their producers, and fimA. A) Scaled expression of C5AR1 by clade. B) Heatmap of the relative abundance of UniRef90 gene families corresponding to canonical gingipains across disease severity groups. C) Correlations between gingipain gene family relative abundances and the relative abundances of their attributed producers. D) Heatmap of the scaled expression of known gingipain target genes by clade. E) Relative abundance of rgpB gingipain across DMN clusters. F) Relative abundance of virulence factor fimA across severity groups.

Finally, we interrogated the relationships between the expression of host genes of interest and the abundance of bacterial metabolic pathways of interest (see Methods). This analysis identified 8 bacterial metabolic pathways with significant associations with three or more of our host genes of interest. Of these, we focused on the pathway with the most associated genes, PWY-5981: CDP-diacylglycerol biosynthesis III (10 associated genes; Figure 5A), and the pathway with the lowest average p-value of associations, PPGPPMET-PWY: ppGpp metabolism (average p-value 0.0014; Figure 5B). All but one of the genes associated with these pathways was significantly differentially expressed between gene expression clades 1 and 2 (*TNIP3* was N.S. adjusted p=0.2; Figure 5C, PWY-5981 genes; Figure 5E, PPGPPMET-PWY genes). The relative abundance of PWY-5981 was significantly elevated in severe disease compared to mild and the relative abundance of PPGPPMET-PWY was significantly elevated in both moderate and severe disease compared to mild (Figure 5D).

**Fig. 5.**
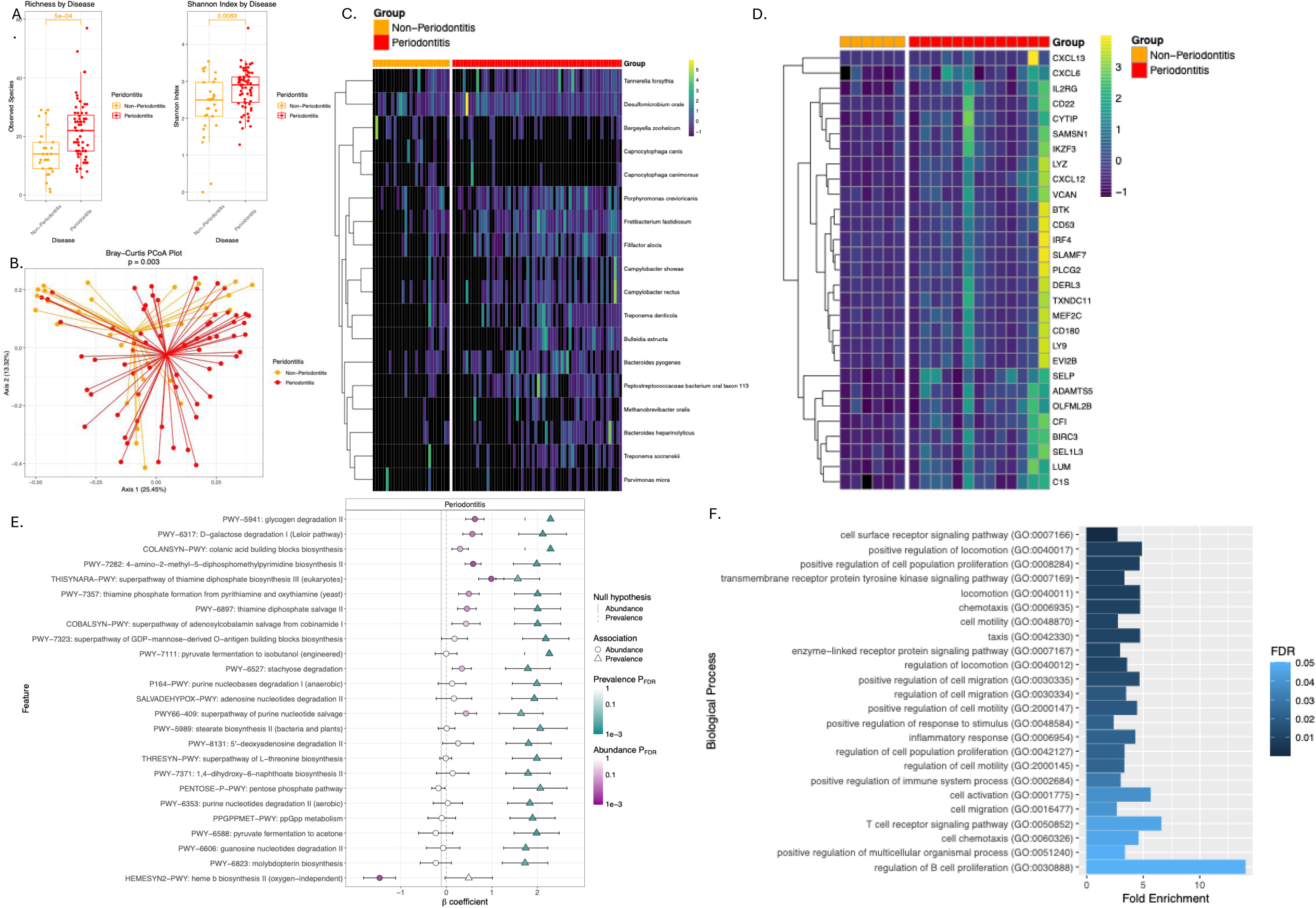
Associations between bacterial metabolic pathways of interest and the expression of host genes of interest. A) Correlation between the relative abundance of PWY-5981: CDP-diacylglycerol biosynthesis III and significantly correlated genes. Y-axis is pathway relative abundance and x-axes is log(relative expression + 1)*1000000 of genes. B) Correlation between the relative abundance of PPGPPMET-PWY: ppGpp metabolism and significantly correlated genes. Y-axis is pathway relative abundance and x-axes is log(relative expression + 1)*1000000 of genes. C) Heatmap of the scaled expression of PWY-5981: CDP-diacylglycerol biosynthesis III pathway-associated genes of interest between clades. D) Relative abundance of pathways of interest across disease severity groups. E) Heatmap of the scaled expression of PPGPPMET-PWY: ppGpp pathway-associated genes of interest between clades. D) Relative abundance of pathways of interest across disease severity groups.

## Discussion

Periodontitis (Perio) is a progressive oral disease characterized by inflammation and degradation of the periodontal apparatus (Martínez-García, 2021). Perio is highly prevalent in domestic canines (Harvey, 1998) and is associated with local and systemic morbidity including loss of teeth, pathological jaw fracture, oronasal fistula formation, cardiovascular disease, diabetes mellitus, among others (Hajishengallis G., 2022).

Response rates to standard-of-care treatments are highly variable, presenting an opportunity for novel therapies and interventions (Kinane, 2017). Furthermore, parallels between canine Perio and human Perio suggest that canines may be promising models for both elucidating common mechanisms involved in Perio and for testing novel treatments (Albuquerque, 2012). These factors make in-depth study of canine Perio a promising area of investigation. In this cross-sectional, retrospective study, we characterize the microbiota and host gene expression in periodontitis in a cohort of companion dogs using shotgun metagenomics and RNA-seq.

We began with an unbiased examination of host gene expression, microbial alpha and beta diversity, and of the bacterial species and metabolic pathways that distinguish Perio from non-Perio. Our findings are broadly consistent with existing literature related to the microbiome of canine Perio in terms of diversity and characteristic species and reveal striking parallels between human and canine Perio in terms of both microbial biomarkers of disease and differences in host transcriptional profiles between Perio and non-Perio. Specifically, the global transcriptional differences we observed in canines with PD was highly consistent with those reported previously in humans (Davanian, 2012) by both gene set enrichment analysis and significantly differentially expressed genes, while the biological processes overrepresented by the differentially expressed genes in our dataset – primarily related to cellular motility (Ivanovski, 2007) and immune response (Zhang M. L., 2024) – are also characteristic of Perio in humans. Furthermore, Perio-associated differences in alpha and beta diversity of the gingival microbiota parallel those reported previously in canines (Davis, 2013).

Additionally, many of the specific species that we found to be elevated in relative abundance in Perio have been linked mechanistically or observationally to the disease in humans, including: *T. forsythia* and *F. alocis* (so-called “red complex” PD-pathogens) (Wirth, 2021), *C. showae* (Macuch, 2000)*, C. rectus* (human and canine Perio) (Özavci, 2019) (Brennan, 2007) (Tamura, 2006), *F. fastidiosum* (Liang, 2025), *M. oralis* (humans and dogs) (Niemiec, The bacteriome of the oral cavity in healthy dogs and dogs with periodontal disease., 2022) (Lepp, 2004), and *T. socranskii* (human and canine Perio) (Takeuchi, 2001) (Nordhoff, 2008). However, several species we observed to be elevated in Perio in our dataset have only been reported in canine Perio and not human Perio, including *B. pyogenes* (Alessandri, 2024), *P. crevioricanis* (Dahlén, 2012), and *D. orale* (Riggio, 2011) (Niemiec, The bacteriome of the oral cavity in healthy dogs and dogs with periodontal disease., 2022). Finally, several of the bacterial metabolic pathways we found to be associated with PD and known to be related to mechanisms of Perio-pathogenesis in humans, including thiamine diphosphate pathways (a factor in *Treponema* virulence) (Bian, 2011), heme biosynthesis (potential deriver of *P. gingivalis* virulence) (Śmiga, 2025) (Guo, 2020), and ppGpp metabolism (important for biofilm formation and *P. gingivalis* virulence) (Kim, Synthesis of ppGpp impacts type IX secretion and biofilm matrix formation in Porphyromonas gingivalis., 2020). Together, these findings both ground this work as being consistent with existing literature on Perio and support the hypothesis that canine PD largely parallels human PD in terms of both host gene expression and, to a lesser extent, changes in the composition of gingival microbiota, the latter of which includes species that appear to be unique to the canine version of the disease.

In the course of our exploratory analysis of host gene expression profiles relative to Perio severity, we observed two distinct clusters or “clades” of samples based on principal component analysis of their overall gene expression. Examining these two clades, we found that the second clade was comprised almost entirely of samples from subjects with moderate disease. Differential expression analysis revealed far more differentially abundant genes between clades than between Perio vs. non-Perio and between the three different severity levels of disease, consistent with a distinct and dramatic host response occurring at a transitional point in disease progression, characteristic of patients going from non-Perio/mild gingivitis to moderate Period with clinical attachment loss. Overrepresentation analysis of biological processes in the highly significantly differentially abundant genes between clades revealed that the abrupt turning point in host response to disease progression is characterized by a shift in immune response, proliferation and adhesion, and cell signaling, consistent with increased inflammation and degradation and disruption of tissue structure/detachment. Additionally, a single pathway was found to be significantly overrepresented in the extremely significantly differentially abundant genes between clades: keratinization, with the expression of component genes observed to be downregulated in clade 2. This is consistent with the destruction and degradation of gingival epithelial cells, which may manifest as gingival tissue detachment and which may enable bacterial invasion and degradation of tissue integrity (Zhang W. Z., 2026). In fact, bacterial invasion of the gingival tissue was confirmed by fluorescence microscopy analysis of samples from this cohort (Supplemental Figure 2).

The observation of an apparent turning point in host response during the intermediate phase of disease progression, in which host gene expression was markedly different compared to early/mild disease and to late/severe disease, motivated us to investigate whether changes in microbiota composition exhibited a parallel three-phase structure with respect to disease severity. Unbiased clustering of samples based on microbiota composition yielded results consistent with this hypothesis: a three-cluster model fit the data better than any other number of clusters, and these clusters were significantly associated with our three disease severity groups. Seven species – six *Porphyromonas* species and *Filifactor alocis* – were the primary drivers distinguishing between the clusters. This is notable because *Porphyromonas* species, particularly *P. gingivalis*, have been implicated as mechanistic drivers of Perio and PD severity via gingipains (e.g. rgpB) and fimbrial proteins (e.g. fimA) (How, 2016) (Kwack, Porphyromonas gulae and canine periodontal disease: current understanding and future directions., 2025). Similarly, *F. alocis* has a well-established association with severe Perio (Aruni A. W., 2015), with proposed mechanisms of virulence including induction of gingival epithelial cell apoptosis (Moffatt, 2011) (Chioma, 2017) and promoting dysbiosis by forming synergistic biofilms with other pathogens (Aruni A. W., 2011). Both *P. gingivalis* (and, to a lesser extent, *P. gulae* in canines (Kwack, Porphyromonas gulae and canine periodontal disease: current understanding and future directions., 2025)) and *F. alocis* have been implicated as “keystone pathogens” in the keystone pathogen hypothesis of Perio (Mishra, 2024). Therefore, a three-phase model of disease progression fits the data well: we observe an abrupt shift in host gene expression as disease progresses to the moderate phase; microbiota composition naturally clusters into three states which parallel disease severity group; and the microbial species driving microbiota clusters have historically been implicated as primary drivers of disease, with pathological effects consistent with both the gene expression changes observed and with the gross tissue morphology that was the basis for our clinical disease severity classifications.

Finally, we turned our attention to previously described mechanisms of virulence to assess whether such mechanistic factors were consistent with our observations. Under the keystone pathogen hypothesis, *Porphyromonas*-derived gingipains cleave complement component C5, generating high levels of C5a and inducing C5a receptor activation, triggering inflammation and additional subversive influences on immune response (Hajishengallis G. D., 2012) (Olsen, 2017).. To assess our hypothesis that this process corresponds to the disease state represented by our clade 2 samples – the abrupt transitional point when disease state shifts away from being mild into moderate severity – we examined *C5AR1* expression across clades, revealing markedly elevated expression in all clade 2 samples except one, which was the one and only clade 2 sample classified as having severe disease. We then visualized other known gingipain responsive genes in host, including *AKT3*, *INSR*, *ICAM1*, *C3AR1*, *TLR2*, *F2R*, *TLR6*, and *TF*, and found them all to be increased in expression in clade 2. We also looked at the relative abundance of *Porphyromonas*-derived gingipain genomic sequences in our metagenomic data and found them all to be elevated in moderate and/or severe disease. To support the attribution of these gingipain sequences to their canonical *Porphyromonas* source species, we showed that the abundance of the source species and the gingipain sequences were highly correlated. We also assessed the relative abundance of metagenomic sequences of the canonical gingipain rgpB across our three unbiased clusters of microbiota samples and found it to be elevated in both the second (corresponding to moderate disease) and third (corresponding to severe disease) clusters, the second cluster having the highest median relative abundance. We also assessed the relative abundance of fimA – another well-established *Porphyromonas*-derived virulence factor (Wang, 2020) – across severity groups, finding it to be significantly elevated in both moderate and severe disease states relative to mild, with the most significant elevation observed in moderate disease. These results support the notion that the abrupt transitional point in disease progression coincides with the action of the canonical keystone pathogen mechanisms.

Lastly, we examined the correlation between the expression of host genes of interest and the abundance of bacterial metabolic pathways, revealing two bacterial pathways that were most associated with host genes implicated in the abrupt shift in host response corresponding to PD progressing beyond moderate severity: CDP-diacylglycerol biosynthesis and ppGpp metabolism. CDP-diacylglycerol, which contributes to virulence by fueling the synthesis of membrane phospholipids and specialized lipids in known PD pathogens *P. gingivalis* (Nichols, A novel phosphoglycerol serine-glycine lipodipeptide of Porphyromonas gingivalis is a TLR2 ligand., 2020) and *T. denticola* (Vences-Guzmán, 2017), was positively correlated with expression of host genes involved in inflammation and platelet activation. These lipids are crucial for bacterial survival in host tissues (Blunsom, 2020), aiding in immune evasion, and promoting periodontal destruction. ppGpp metabolism, which is an indicator of stress response in bacteria (Liu, 2015) and regulator of proliferation and biofilm formation in *Porphyrnomonas* (Kim, Synthesis of ppGpp impacts type IX secretion and biofilm matrix formation in Porphyromonas gingivalis., 2020), was negatively correlated with *F2R*, a known gingipains-responsive gene (Lourbakos, 2001) (Nylander, 2008); and with *DCBLD2* and *TNIP3*, genes involved in wound healing (Schmoker, 2019) and inflammation (Wullaert, 2007), respectively. These findings are consistent with two potential mechanisms of virulence under the keystone pathogen hypothesis: A) CDP-diacylglycerol increases in abundance during the abrupt stage in disease progression, enhancing the pathogenic virulence which drives the corresponding abrupt host response, and B) ppGpp metabolism – a master regulator of stress-related bacterial metabolism – parallels the abrupt host response as pathogens adapt to the more severe disease state.

Notably, the cross-sectional, observational nature of this study constitutes a significant limitation: we cannot directly evaluate our mechanistic hypotheses, or our hypothesis relating to disease progression over time within individuals. However, our observations consistent with these hypothesize suggest appealing targets for future work. Specifically, we suggest a longitudinal study focused on monitoring a cohort of individuals over time to validate our hypothesis of an abrupt change in microbiota and host response at the transitional point when disease progresses from mild to moderate severity. Additionally, we suggest that the mechanisms driving this abrupt transitional period can be studied in canines both for their potential value to veterinary medicine and for their potential translation value in human PD. Finally, the high proportions of canine genomic sequences in our shotgun metagenomic samples impose limits on the kinds of analyses we can do, including our ability to characterize virulence and antimicrobial resistance factors. As such, we recommend deeper sequencing for any subsequent metagenomic studies of similar sample types.

The identification of a distinct, abrupt molecular and microbial shift marking the transition from mild to moderate periodontitis has meaningful implications for both veterinary and human oral medicine. This transitional phase, driven by the proliferation of keystone pathogens like *Porphyromonas spp.* and *F. alocis* and characterized by a sudden downregulation of host epithelial keratinization and an upregulation of destructive inflammatory pathways, represents a critical clinical window. Interventions targeted at this specific juncture – perhaps by inhibiting the canonical virulence factors (such as gingipains) or disrupting the associated bacterial metabolic pathways (like CDP-diacylglycerol or ppGpp metabolism) identified here – could prevent the onset of irreversible clinical attachment loss. Furthermore, by confirming that the keystone pathogen hypothesis robustly applies to the canine disease model, these findings validate the domestic canine not merely as a proxy, but as a highly relevant translational platform for elucidating the nuanced host-microbiome crosstalk that dictates disease progression.

## Conclusions

This study provides a comprehensive metagenomic and transcriptomic characterization of canine periodontitis, revealing striking etiopathogenic parallels with the human disease. We demonstrate that the progression of periodontitis is not a simple linear degradation, but rather features a pivotal, pathogen-driven inflection point during the transition to moderate disease, characterized by abrupt topological shifts in the microbiome and a corresponding, dramatic structural and immunological host response. By mapping the interplay between host gene expression, specific microbial shifts, and the mechanistic virulence factors of keystone pathogens, this work advances our fundamental understanding of veterinary periodontal disease. Ultimately, these insights underscore the value of the companion dog as a robust model for studying periodontal pathogenesis, offering promising new diagnostic biomarkers and therapeutic targets for mitigating this highly prevalent disease across species.

## List of abbreviations

PD: Periodontal Disease
Perio: Periodontitis
DMN: Dirichlmet Multinomal

## Declarations

### Ethics approval and consent to participate

All procedures involving companion dogs were approved by Cornell University’s Institutional Animal Care and Use Committee (IACUC) (Protocols #2015-0117 and #2015-0219). Participation was strictly voluntary, and written informed consent was obtained from all dog owners prior to the collection of microbial samples and gingival tissues.

### Consent for publication

Not applicable.

### Availability of data and material

The datasets generated and analyzed during the current study (shotgun metagenomic sequences and RNA-seq reads) are available in the NCBI Sequence Read Archive (SRA) repository under BioProject accession number PRJNA1442231.

### Author contributions

AG, NF, JKG and SP conceived and designed the study. NF, MJB, JKG, AMV, NT and SP performed experiments and/or collected samples. AG, JKG, and SP analyzed the metagenomic and/or transcriptomic data and performed statistical modeling. AG and SP drafted the manuscript. All authors provided critical revisions to the text and approved the final manuscript.

## Acknowledgements

This research was supported by a grant from the AKC Canine Health Foundation (Grant No. 02809). Metagenomic sequencing was conducted by the Weill Cornell Medicine Microbiome Core. Gene expression profiling was conducted by the Transcriptional Regulation and Expression Facility at Cornell University. Experimental sample collection and processing, as well as client-owned dog enrollment were kindly facilitated by the Cornell University Hospital for Animals Clinical Trials group, led by Ms. Carol E. Frederick, LVT, VTS (ECC) and Ms. Andrea L. King, LVT; by Cornell’s Innovation laboratory; and by the Dentistry and Oral Surgery technical staff and clinicians. Cryopreserved tissue samples and associated phenotypic data were provided by the Cornell Veterinary Biobank, a resource built with the support of NIH grant R24 GM082910 and the Cornell University College of Veterinary Medicine.

## Legends

Supp. Fig 1. GSEA Enrichment score and Ranked list metric plots of Gene Set Enrichment Analysis of all human homolog genes against the Davanian 2012 gene set, based on the comparison between Perio and Non-Perio.

Supp. Fig 2. Localization of Porphyromonas and Tannerella spp. within gingival tissue from periodontitis-affected canine patients. (A) DAPI-stained image of gingival tissue from patient 7268 AA2457, with boxed region indicating the area shown in (B). Scale bar = 500 µm. (B) Higher-magnification view of the boxed region in (A), demonstrates the presence of Tannerella spp., localized to host cell nuclei. Scale bar = 20 µm. (C) DAPI-stained image of gingival tissue from patient 29839 AA2334, with boxed region indicating the area shown in (D). Scale bar = 1 mm. (D) High-magnification field of view of the boxed region in (C), demonstrates the presence of Porphyromonas spp., localized at the periphery of host cell nuclei. Scale bar = 20 µm.

Supp. Table 1. Complete list of significantly differentially abundant/prevalent microbial pathways between Perio and Non-Perio and corresponding statistics.

## Supplemental data

Supplemental Figure 1 - Supp_Fig_1_Davanian_Pr_vs_non-Pr_UP_1.png

Supplemental Figure 2 - Supplementary_Figure_2.docx

Supplemental Table 1 - Supplementary_Table_1.tsv

