## Supplementary figures and images for "Metagenomic and transcriptomic signatures of periodontitis in companion dogs"

### Supplementary Figure 1

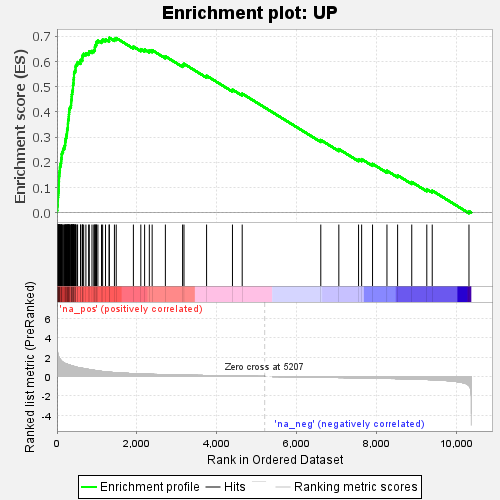
